# Vacuolar invertase knockout enhances drought tolerance in potato plants

**DOI:** 10.64898/2025.12.01.691554

**Authors:** Marina Roitman, Paula Teper-Bamnolker, Adi Doron-Faigenboim, Noga Sikron, Aaron Fait, Ondrej Vrobel, Petr Tarkowski, Menachem Moshelion, Samuel Bocobza, Dani Eshel

## Abstract

Drought stress is one of the most critical abiotic constraints limiting crop productivity worldwide, exacerbated by ongoing climate change and increasingly frequent extreme weather events. Stomatal regulation and osmoprotective sugar accumulation are critical adaptive mechanisms for plant survival under drought stress. Here, we characterize the enhanced drought resilience observed in CRISPR/Cas9-mediated potato, mutants in their vacuolar invertase gene (*StVInv*). Knockout plants exhibited improved performance under progressive drought stress and during rewatering drought, maintaining higher stomatal conductance, elevated transpiration rates, and superior photosynthetic efficiency compared to wild-type (WT) plants. These improved performance under similar transpiration rate led to higher agronomic water-use efficiency (AWUE) in *stvinv* plants resulting in greater biomass production despite reduced water availability. Metabolomic profiling revealed distinct adaptive strategies; *stvinv* plants preferentially accumulated galactinol and raffinose, indicating enhanced raffinose family oligosaccharide (RFO) metabolism. Furthermore, *stvinv* plants displayed lower levels of abscisic acid (ABA) and its catabolites under drought, suggesting a moderated ABA response facilitating a more risk-taking growth strategy that supports sustained growth and physiological stability. Our findings identify targeted metabolic and hormonal adjustments underlying drought resilience in potato plant, offering promising strategies for enhancing crop performance under water-limited conditions.

## Introduction

Abiotic stresses impair plant growth and development, reduce productivity, and limit plant species’ geographical distribution [1]. Plants’ survival depends on their timely and pertinent responses to changes in growth conditions, the severity and duration of stress conditions, and the capacity to adapt quickly to changing energy equations [2]. Plants cope with abiotic stress by differentially regulating various genes at the transcriptional level [reviewed by 3]. These stress-responsive genes play a role in stress-signal transduction and gene-expression regulation; the products from these genes protect plant cells from stress-related damage and maintain cell viability [4]. The antioxidant defense machinery protects plants against oxidative stress damage [5]. Plants possess efficient enzymatic and non-enzymatic antioxidant defense systems. These systems work in concert to control the cascades of uncontrolled oxidation and protect plant cells from oxidative damage by scavenging of reactive oxygen species (ROS) [6, 7].

Osmotic adjustment is an effective component of stress tolerance. Accumulation of osmoprotectants (proline, glycine betaine, gamma-aminobutyric acid, and sugars) is a typical response to abiotic stresses observed in different plant systems [8]. Modulating the expression of genes related to the production and accumulation of compatible solutes helps plants to tolerate osmotic stress by maintaining water potential and protecting cellular organelles and essential proteins [9]. Soluble sugars are important osmoprotectants that play a significant role in cellular osmotic adjustment by protecting cell structures exposed to environmental stress [10–13]. Sucrose and sucrosyl oligosaccharides, including fructans and RFOs, accumulate in the vacuoles under oxidative stress as a consequence of abiotic stress, protecting the tonoplast against ROS-mediated damage [14]. During stress, sucrose and RFOs can be transported from the vacuole to the apoplast as tonoplast exocytosis vesicles for stabilization of cell membrane [15, 16].

Sucrose and its cleavage products glucose and fructose are central molecules for cellular biosynthesis and signal transduction throughout the plant life cycle [17]. Sucrose may serve as a protectant against cold stress by acting as a signal or as an osmoprotectant (cryoprotective) molecule [16, 18]. *In-vitro* studies have shown that the ID_50_ value required to inhibit OH^•−^ catalyzed hydroxylation by sucrose is similar to that of glutathione antioxidant [19]. In agreement with this, PpINH1, an invertase inhibitor in peach, maintains high sucrose levels, improves membrane stability during cold storage, and enhances resistance to chilling injury [20, 21].

Other essential water-soluble carbohydrates derived from sucrose include the RFOs (α-galactosyl extensions of sucrose) produced by galactinol synthase (GolS) and raffinose synthase (RafS) using sucrose and myo-inositol as substrates. RFOs are proposed to play an essential role in protecting plants from oxidative stress, as they accumulate under stressful conditions [19, 22–24]. In addition to functioning as osmolytes, RFOs also participate in carbon storage, in stress signaling pathways, in membrane protection and as antioxidants against ROS in different cell compartments [16, 19, 25–27].

Accordingly, inhibition of the *VACUOLAR INVERTASE* gene (VInv) in *Arabidopsis thaliana* or *Solanum tuberosum* promoted accumulation of raffinose, increasing the cold tolerance of transgenic plants [28, 29]. Generally, rice (*Oryza sativa*) and *Arabidopsis* do not accumulate large quantities of RFOs in tissues under optimal conditions. However, RafS accumulation was observed under stressed conditions, such as extreme temperature [30, 31]. Although there is evidence of a correlation between GolS activity and RFOs content, the concentration of the initial substrates (myo-inositol and sucrose) is the key for RFOs accumulation at least in seeds and tubers [29, 32–34]. Sucrosyl oligosaccharides and the enzymes associated with their metabolism may interact indirectly with ROS-signaling pathways [27, 35]. In addition, RFOs and galactinol have been proposed to play important roles in oxidative-stress protection in plants during stress acclimation [15, 19, 36]. All these previous studies show that the production of sugars is beneficial for plant survival during abiotic stress.

A previous review suggests that ROS signaling interacts with ABA, Ca²⁺ fluxes, and sugar sensing, and may function both upstream and downstream of ABA-dependent pathways during drought stress [37]. This has implied that RFOs may play a pivotal role in the initial adaptation of plants to water-deficient conditions, potentially by controlling both ROS signaling and metabolism [16, 38]. Plant’s responses to drought stress are closely linked to the hormone ABA [39]. Drought-related gene expressions are believed to be governed through both ABA-dependent and ABA-independent mechanisms [40]. Numerous elements of the ABA signaling pathway and the control of gene expression downstream of ABA signaling have been thoroughly documented, but the ABA-independent pathway remains elusive and not well understood [41]. When plants experience stress, the production of ABA from scratch relies on the activation of the *NCED3* gene [42]. This gene codes for a 9-cis-epoxycarotenoid dioxygenase enzyme, which plays a crucial role in the primary step of ABA biosynthesis. In situations where roots face water scarcity, a hydraulic signal is generated, leading to a swift transmission of a water deficiency signal from the roots to the leaves of the plant. This, in turn, initiates the synthesis of ABA in the leaves and the closing of stomata [43].

In the study of the ABA biosynthesis mutant *nced3* subjected to dehydration stress, researchers observed that proline (an amino acid involved in plant stress response), branched-chain amino acids (BCAAs), and gamma-aminobutyric acid (GABA) accumulate later and in lower quantities compared to sugars, and this pattern was influenced by the presence of ABA. On the other hand, RFOs and the antioxidant ascorbate increased independently of ABA [44]. These results suggest that the ABA-independent response to drought likely occurs earlier than the ABA-dependent response [38].

Drought stress poses a major challenge to the production of potatoes worldwide. Climate change is predicted to further aggravate this challenge by intensifying potato crop exposure to increased drought severity and frequency [45]. Drought stress was reported to trigger the accumulation of soluble sugars in sink leaves of potato plants, reduction in tuber number and yield and affect carbon partitioning in the whole plant [45, 46]. Although the application of exogenous ABA has confirmed the ABA-induced expression of stress-related genes, several drought-induced genes are insensitive to exogenous ABA application [45]. The Mechanism of ABA-independent drought response pathway has not been described, to our knowledge, in any plant species. We are proposing to elucidate an ABA-independent pathway for drought tolerance in potato. Although potato has not been a classical model for molecular biology and genomics research until recently [47, 48], it provides an ideal “case study” to dissect the ABA-independent drought response pathway and determine its effect on agricultural crop performance.

The potato tuber is a swollen underground stem formed by swelling of the subapical underground stolons [49]. Exposing the potato tuber to chilling during field-growth or postharvest storage triggers cold-induced sweetening (CIS), characterized by the accumulation of hexoses, such as glucose and fructose, mainly in the tuber parenchyma [50, 51]. Accordingly, potato tubers have been shown to produce free radicals under low-temperature conditions [52]. Sucrose is cleaved to hexoses by two main enzymes: sucrose synthase (EC 2.4.1.13) and invertase (EC 3.2.1.26). Sucrose synthase catalyzes the reversible conversion of sucrose to uridine diphosphate-glucose and fructose. Invertases irreversibly split sucrose into glucose and fructose [reviewed by 53]. Sucrose hydrolysis by *S. tuberosum* vacuolar invertase (StVInv) has been reported to be the main pathway involved in potato CIS [54–56]. In a previous study, we demonstrated that potato plants with CRISPR knocked-out *StVInv* genes (*StVInv*) exhibit enhanced tolerance to combined cold and drought stress, associated with increased RFO accumulation in tubers [29]. Here, we dissected the specific contributions of the *StVInv* gene to drought tolerance. Using comprehensive physiological phenotyping, including continuous transpiration and stomatal conductance measurements, alongside detailed metabolomic analyses, we characterized the unique drought adaptation strategies of *stvinv* plants. Our findings revealed that these knockout plants exhibit enhanced drought resilience through sustained stomatal function, moderated ABA signaling, and a distinct metabolic shift toward increased osmoprotective sugar accumulation. These findings underscore the central role of sucrose metabolism in plant drought adaptation, providing valuable targets for breeding drought-resilient crops.

## Results

### Enhanced drought tolerance in *stvinv* plants

We previously demonstrated the enhanced tolerance of two independent CRISPR/Cas9 knockout lines of *StVInv*, referred to as *stvinv-7* and *stvinv-8* potato plants to combined cold and drought stress during early seedling stages (Teper-Bamnolker et al., 2023). To isolate the specific effects of drought stress and elucidate the underlying mechanisms of drought response in *stvinv* plants, we used one-month-old seedlings. Plants were grown under controlled conditions, exposed to prolonged terminal drought followed by rehydration to assess recovery, and their response was visually scored on a qualitative phenotypic scale from low to high vigor. In all tested lines, the phenotype score began to decline approximately 14 days after irrigation was stopped (Fig. 1A). Throughout the drought period, *stvinv* plants exhibited significantly greater tolerance compared to WT plants, as evidenced by higher phenotypic scores (Fig. 1A, B). Chlorophyll content was higher in *stvinv* plants during the irrigation phase and the early days of drought, as well as during the initial recovery period, suggesting enhanced photosynthetic efficiency compared to WT plants (Fig. 1C). *stvinv* plants maintained elevated leaf stomatal conductance relative to WT throughout all phases—irrigation, drought, and recovery. Notably, both *stvinv-7* and *stvinv-8* exhibited significantly higher stomatal conductance during severe drought and recovery (Fig. 1D). This enhanced gas exchange capacity likely facilitated sustained photosynthetic activity, improved water-use efficiency, and expedited recovery following drought stress.

**Fig. 1.**
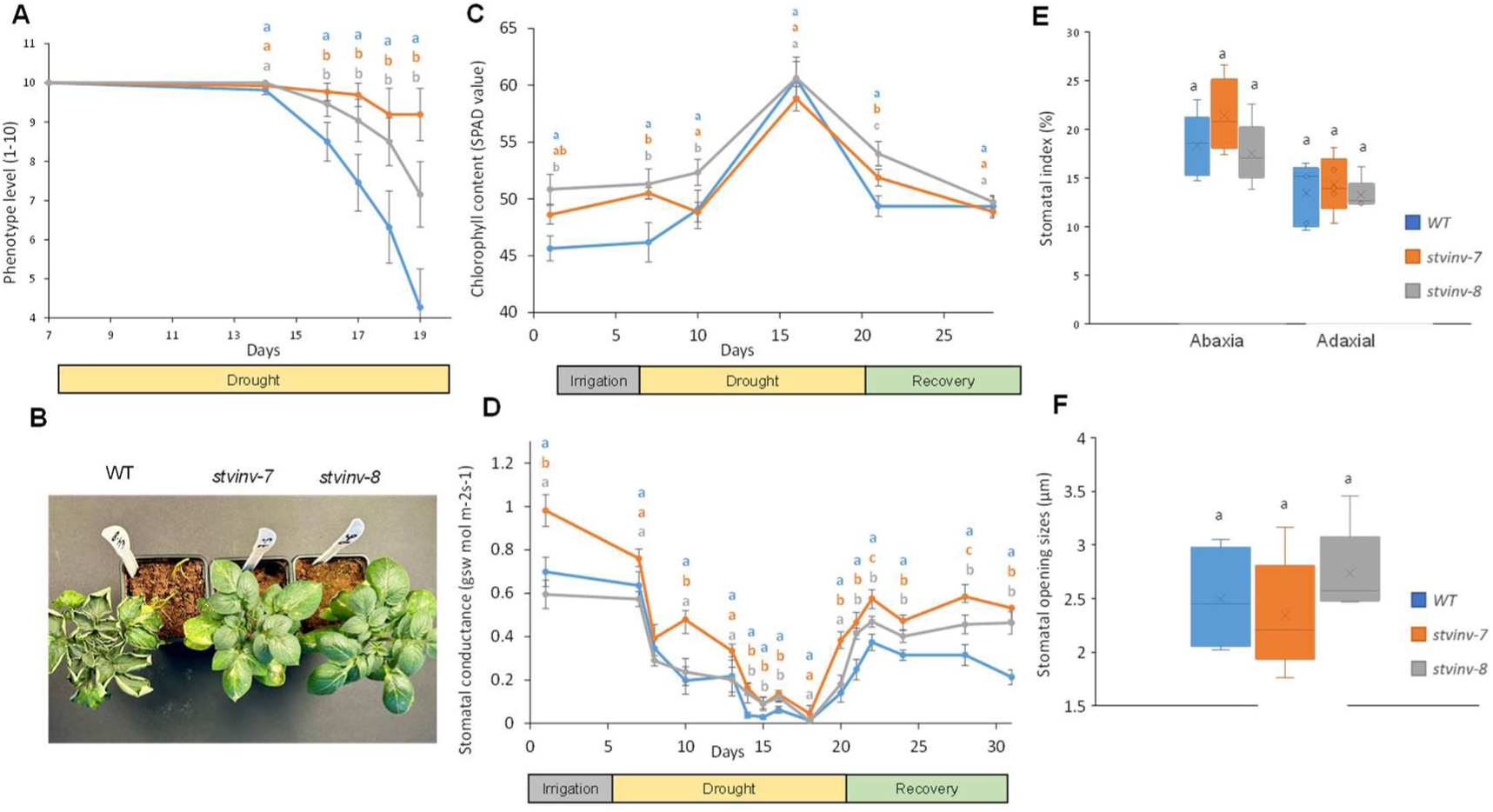
*stvinv* plants show resilient phenotypes under drought conditions with higher chlorophyll content and stomatal conductance. Potato plants of wild type (WT) and *stvinv* lines (*stvinv-7*, and *stvinv-8*) were grown under controlled conditions with a 16-hour photoperiod at 23°C and then subjected to drought treatment (days 15-19), and irrigated again until full recovery. **A**, Phenotype visual score during drought treatment. **B**, representative plants at the end of the drought treatment. **C**, Changes in chlorophyll content; **D**, Leaf stomatal conductance. Different letters above data points indicate statistically significant differences between genotypes (p < 0.05). **E,** Stomatal index calculated as the ratio of stomatal number to total epidermal cells, with stomatal density (number of stomata per mm²) quantified by counting stomata in calibrated microscopic fields. **F**, Stomatal opening size (aperture width) measured from epidermal peels using light microscopy and analyzed with ImageJ software.

### Drought tolerance of *stvinv* plants is independent of stomatal density

To assess whether the different drought response in *stvinv* plants is associated with stomatal traits, stomatal density and aperture width were analyzed. Young, fully expanded leaves were tested at mid-morning under controlled long-day photoperiod conditions. Imprints of abaxial and adaxial leaf surfaces were made following the protocol described by Geisler and Sack [57]. Microscopic analysis revealed no significant differences in stomatal density or indices between *stvinv* and WT leaves across all examined surfaces, abaxial and adaxial (Fig. 1E, F). Measurements showed no significant differences between WT and *stvinv* leaves on both abaxial and adaxial leaf surfaces (Fig. 1F). These findings indicate that the enhanced drought tolerance observed in *stvinv* plants is not attributable to changes in stomatal density or aperture size.

### *stvinv* plants demonstrate higher transpiration rate

To elucidate the physiological water relations mechanisms underpinning drought response, whole-plant continuous transpiration measurements were conducted using the high-throughput telemetric, gravimetric-based phenotyping system (Fig. 2A; Plantarray 3.0, Plant-DiTech, Israel; [58]). This method provided real-time, high-resolution data on water dynamics, facilitating a detailed analysis of transpiration behavior during drought stress and recovery in *stvinv* plants. Two-month-old plants were subjected to a controlled irrigation regime consisting of three phases: an initial well-watered phase (days 1–12) with excessive irrigation, a standardize drought phase (days 13–27) during which daily irrigation was gradually reduced to 80% of each plant’s transpiration from the previous day (enabling similar drought stress to all plants), and a recovery phase (days 28–34) with resumed full irrigation.

**Fig. 2.**
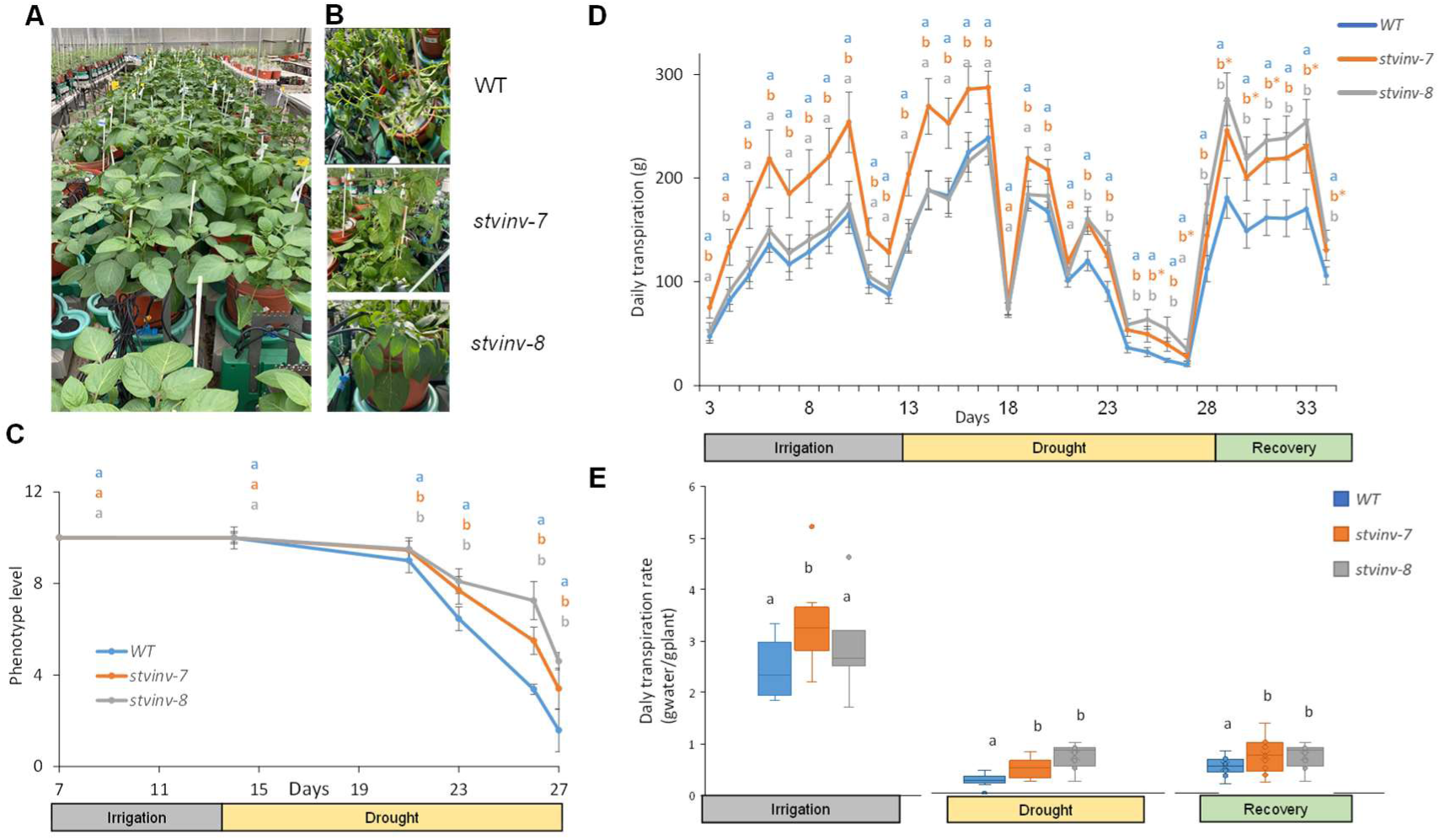
*stvinv* plants demonstrate enhanced drought tolerance. **A,** Experimental design using PlantArray 3.0. Plants were placed on load cells for continuous monitoring of transpiration and conductance. Drought stress was initiated on day 13 for a subset, followed by rehydration on day 27; controls remained well-irrigated throughout. **B,** Representative images of plants on day 25 (corresponding to day 13 of drought), highlighting genotype-specific responses. **C**, Phenotypic assessment of plant vigor during drought and recovery (scale 1–10; 1 = severe wilting, 10 = fully vigorous). **D,** Whole-plant daily transpiration rates across irrigation (days 1–12), drought (days 13–27), and recovery (days 28–34). **E,** Normalized whole-plant daily transpiration rates (per unit biomass). Different letters above data points indicate statistically significant differences between genotypes (p < 0.05).

During this assay, WT plants began showing visible wilting symptoms on day 21, eight days after irrigation was stopped, and exhibited severe wilting by day 27. In contrast, *stvinv* lines maintained higher phenotypic scores during the late drought phase (days 21–27), with significantly milder stress symptoms (Fig. 2B and C). Following rehydration on day 27, both *stvinv* and WT plants start to recover within 24 h (Fig. 2D).

Also, during the well-watered phase (days 1–12), plants exhibited similar transpiration rates, with *stvinv-7* plants generally trending higher (Fig. 2D). Under drought conditions (days 13–27), WT plants experienced a significant decline in transpiration rates, particularly between days 19-26. In contrast, *stvinv* lines maintained substantially higher transpiration rates throughout this phase (Fig. 2D). During recovery from drought (days 28–34), both *stvinv* lines, particularly *stvinv-8*, demonstrated significantly higher transpiration rates than WT plants, highlighting their rapid resilience (water-use resumption post-rehydration; Fig. 2D). Midday whole-canopy conductance measurements (10:00–15:00) confirmed that *stvinv* plants consistently maintained higher conductance levels under drought stress compared to WT plants (Fig. S1). This higher gas exchange capacity reflects their ability to regulate water loss while sustaining physiological functions. Additionally, normalized transpiration rates (calculated as transpiration relative to biomass) further underscored the drought resilience of *stvinv* plants, with significantly higher rates during both the drought and recovery phases (Fig. 2E).

Whole-canopy transpiration rates, normalized to biomass (E), highlight consistently higher transpiration levels in *stvinv* plants compared to WT plants under drought conditions (Fig. 3A). On peak drought days (e.g., day 23), E measured during daylight hours (06:00–18:00) provided insights into stomatal dynamics. Normalizing the transpiration rate to VPD, revealed that *stvinv* plants exhibited consistently elevated midday conductance (Fig. 3C) and transpiration (Fig. 3B), without temporal variation, indicating a stable, robust drought adaptation strategy. This stable, ‘risk taking’, transpiration behavior suggests more carbon fixation and evaporative cooling, contributing to the enhanced drought resilience of *stvinv* plants.

**Fig. 3.**
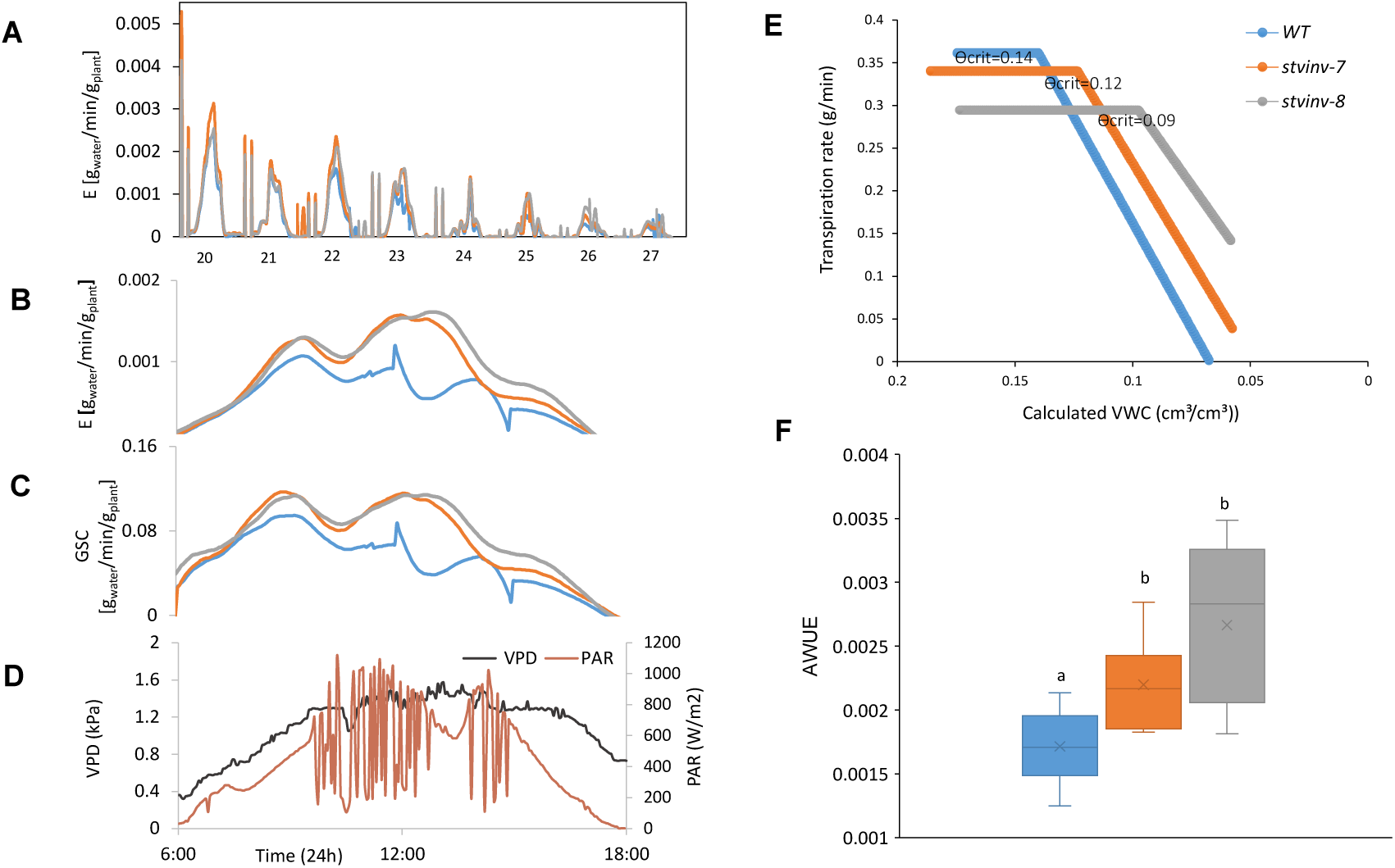
Physiological responses of WT and *stvinv* plants under drought conditions. **A**, Whole-canopy normalized transpiration rates (E) of WT, *stvinv-7*, and *stvinv-8* monitored continuously during the drought period (days 20–27) between 06:00 and 18:00, under natural photosynthetically active radiation (PAR) and vapor pressure deficit (VPD). **B,** Daily variation in environmental conditions on a representative drought day (day 23), showing PAR (red) and VPD (black). **C,** Whole-canopy stomatal conductance (GS_C_) measured continuously on day 23. **D,** Whole-canopy transpiration rates (E) on day 23. **E,** Critical soil water content (Θ_crit), defined as the volumetric water content (VWC) at which transpiration is restricted. **F,** Agronomic water-use efficiency (AWUE), calculated as shoot dry weight per total water transpired. Boxplots show medians, interquartile ranges, and individual values. Different letters indicate statistically significant differences (p < 0.05). Error bars represent ±SE; n = 10–13 per group.

### *stvinv* plants present a better water-use efficiency

Soil moisture thresholds (Θ_crit) at which plants restricted transpiration were determined to evaluate the whole plant water balance regulation. WT plants began restricting transpiration at higher soil moisture levels (Θ_crit = 0.14 cm³/cm³) compared to *stvinv* plants (Θ_crit = 0.12 cm³/cm³ for *stvinv*-*7* and 0.09 cm³/cm³ for *stvinv-8*) (Fig. 3E). This gradual response to soil drying in *stvinv* plants enabled sustained transpiration, improving their ability to utilize available water efficiently during prolonged drought.

Both *stvinv-7* and *stvinv-8* lines demonstrated significantly higher AWUE compared to WT plants, as calculated by the ratio of shoot dry biomass to total transpired water (Fig. 3F). *stvinv-8* exhibited the highest AWUE values, indicating superior biomass production per unit of water consumed. These results suggest that *stvinv* plants utilize a better water use efficiently under drought conditions.

### *stvinv* plants exhibit enhanced biomass production following drought stress

To assess whether the drought tolerance of *stvinv* plants translates into improved growth and productivity, shoot, root, and tuber parameters were evaluated at the conclusion of the 35-day experiment. *stvinv* plants, particularly *stvinv-8*, demonstrated significantly higher shoot dry weight compared to WT plants, reflecting enhanced biomass accumulation under stress conditions (Fig. 4A). Root length measurements showed no significant differences among genotypes, indicating that drought tolerance in *stvinv* plants is not linked to root elongation (Fig. 4B). Additionally, the *stvinv-7* line produced a slightly higher number of tubers compared to WT and *stvinv-8* (Fig. 4C). These findings demonstrate that *stvinv* plants not only exhibit drought tolerance but also effectively convert this resilience into enhanced shoot biomass, underscoring their potential for improved growth in water-deficient conditions.

**Fig. 4.**
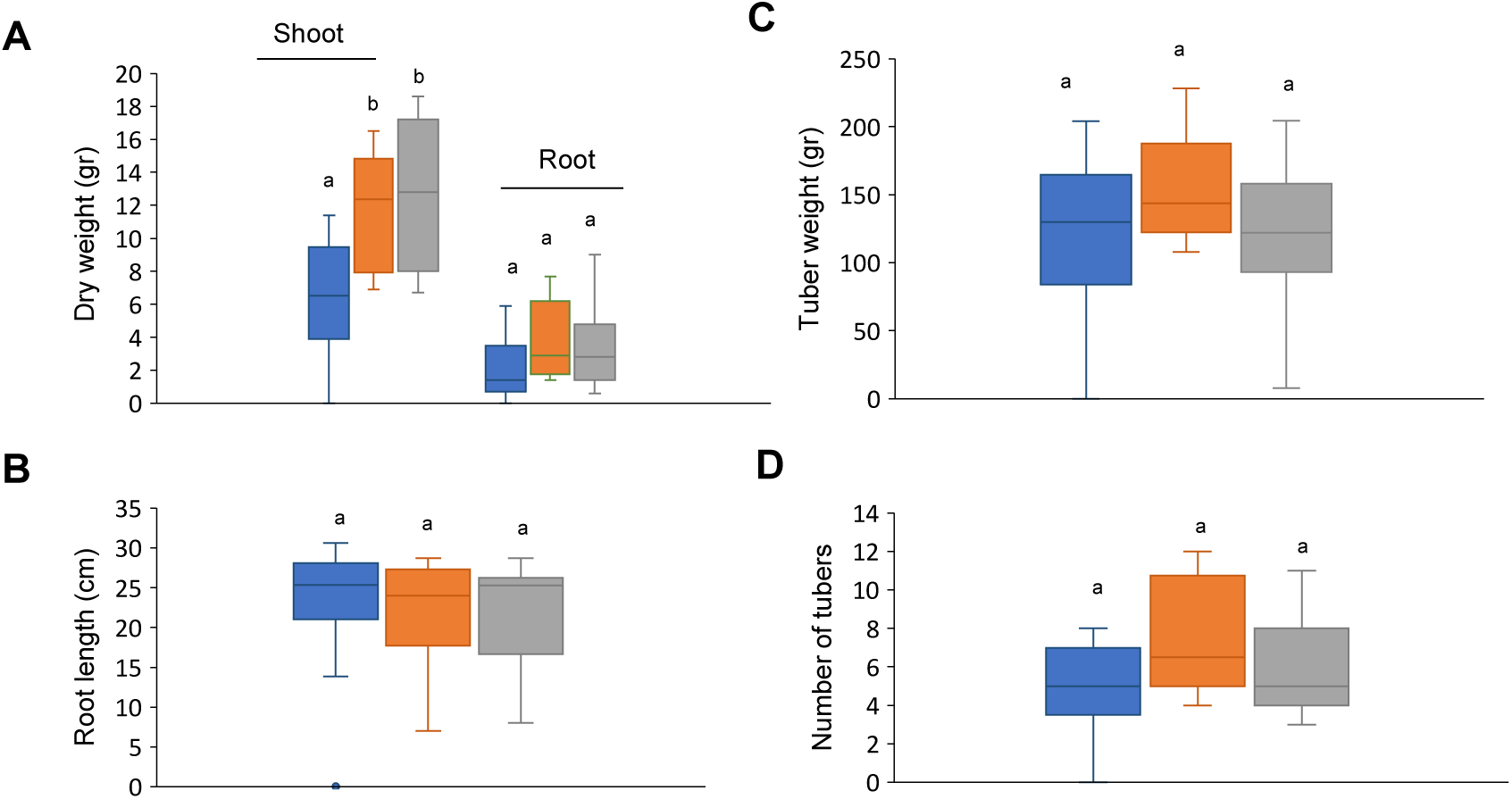
Enhanced biomass and tuber yield in *stvinv-7* and *stvinv-8* plants under drought conditions. **A,** Shoot dry weight, **B,** root dry weight, **C,** number of tubers, **D,** total tuber weight, and **E,** tuber size were measured at harvest (day 34) and are presented as box-and-whisker plots. Different letters indicate statistically significant differences between genotypes (Student’s t-test, p < 0.05). n = 10–13 plants per group.

### *stvinv* plants display enhanced osmoprotection and lower stress perception under drought

To investigate the metabolic adaptations associated with the *StVInv* knockout and its potential role in drought tolerance, we performed a comprehensive metabolomic profiling. Fully expanded young leaves from the third node below the apical bud were sampled at five time points: baseline (T0, following 10 days of irrigation); mild drought (T1, T2; after 7 and 9 days without irrigation); progressive drought stress (T3, T4; after 12 and 14 days without irrigation); and recovery (T5, one day post-rewatering). Samples were analyzed using gas chromatography mass spectrometry (GC-MS) for untargeted metabolomic profiling. Metabolite analysis revealed that WT plants displayed greater level of homoserine and shikimic acid, which are markers of intensified amino acid and secondary metabolism, following drought stress (Fig. 5A). *stvinv* plants consistently accumulated significantly higher levels of osmoprotective sugars such as galactinol and raffinose (Fig. 5B), indicating the distinct stress response strategies between genotypes.

**Fig. 5.**
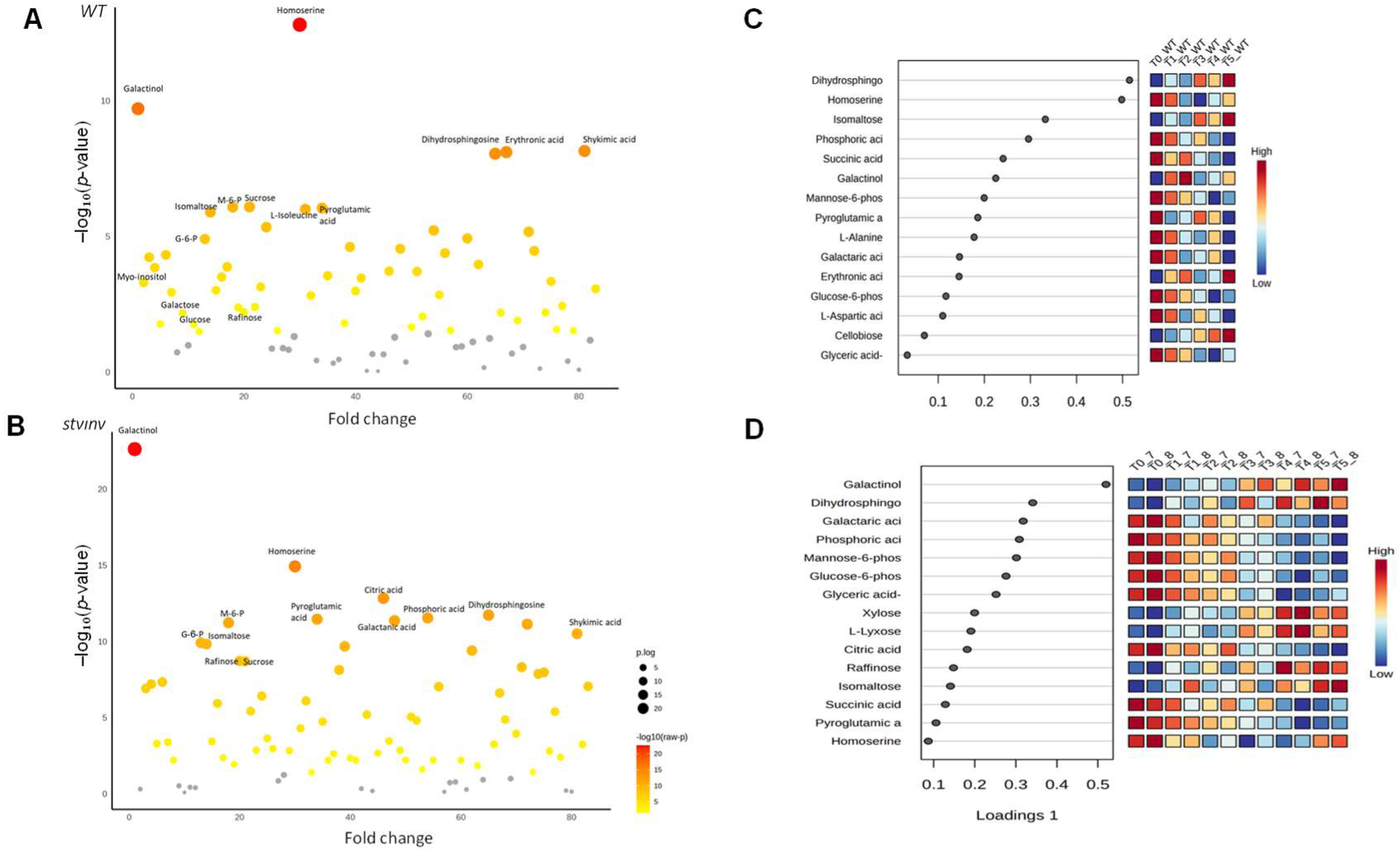
Divergent metabolic responses to drought stress in WT and *stvinv* plants. **A-B,** Volcano plots showing drought-induced changes in metabolite abundance for WT (A) and *stvinv* plants (B). The x-axis indicates fold change; the y-axis shows –log₁₀(*p*-value). Larger bubbles and warmer colors (yellow to red) represent metabolites with greater statistical significance. Only metabolites with *p* < 0.05 (ANOVA with post hoc tests) are shown. **C-D,** Sparse PLS-DA of leaf metabolite profiles across drought stages (T0, irrigation; T1–T2, early drought; T3, mid-drought; T4–T5, severe drought/recovery) in WT (C) and stvinv plants (D). For each genotype, the dot plot shows the loadings of metabolites on component 1, indicating their contribution to separation among drought stages, and the adjacent heatmap shows their scaled abundance across time points and biological replicates, illustrating a shift from amino-acid and sphingolipid associated markers in WT to persistent enrichment of galactinol, raffinose and related sugars in stvinv.

Pathway enrichment analysis confirmed this divergence: WT plants exhibited broad activation of stress-related pathways including glycine, serine, threonine and galactose metabolism and amino acid biosynthesis (Fig. S2), whereas *stvinv* plants showed similar pattern but in higher levels of galactose and sucrose metabolism and unique glyoxylate/dicarboxylate higher metabolism (Fig. S2). This metabolic adjustment in *stvinv* lines appears more energy-efficient, prioritizing carbon conservation and osmoprotection over protein turnover. Notably, this efficiency is further enhanced by increased activation of glyoxylate and dicarboxylate metabolism, facilitating carbon conservation and maintaining energy balance under drought stress. Additionally, elevated citric acid levels in *stvinv* plants point to a more efficient TCA cycle, providing sufficient ATP to support cellular functions during water deficit. Correlation heatmaps further highlighted these differences, showing a strong positive cluster of sucrose, galactinol, and raffinose in *stvinv* plants (Fig. S3A), reflecting a coordinated sugar-based response. WT plants, in contrast, showed more diffuse correlation networks involving amino acids and secondary metabolites (Fig. S3B), consistent with a broader metabolic adjustment under stress. Sparse PLS-DA analysis WT plants were characterized by early-stage peaks in homoserine and dihydrosphingosine, suggesting a transient stress-reactive response (Fig. 5C). In *stvinv* lines, galactinol, raffinose, and glucose-6-phosphate levels were higher than WT the drought and recovery phases (Fig. 5D). By reducing reliance on energy-intensive pathways and enhancing osmoprotective sugar accumulation and carbon recycling, the *stvinv* plants minimized stress perception and maintained physiological homeostasis more effectively than their WT counterparts. Together, these findings indicate that *stvinv* plants adopt a focused, sugar-centered metabolic strategy to mitigate drought stress.

### Drought tolerance of *stvinv* is associated with elevated RFO metabolism in leaves

Our previous work demonstrated that *stvinv* plants showed increased expression of RFO biosynthetic genes followed by higher levels of RFOs in potato tuber parenchyma in response to cold stress [29]. Given this finding, we hypothesized that the enhanced drought tolerance observed in *stvinv* plants might similarly involve RFO metabolism. Metabolomic profiling revealed that all genotypes exhibited a general trend of reduced glucose-6-phosphate and sucrose levels, alongside increased galactose, galactinol, and raffinose levels during drought and recovery phases (Fig. 6). However, *stvinv* plants showed, during drought and recovery from drought, higher levels of sucrose, galactinol and raffinose, and lower levels of myo-inositol compared to WT plants (Fig. 6). These results suggest that *stvinv* plants may metabolize RFO more effectively, contributing to their ability to maintain physiological and biochemical homeostasis under drought conditions.

**Fig. 6.**
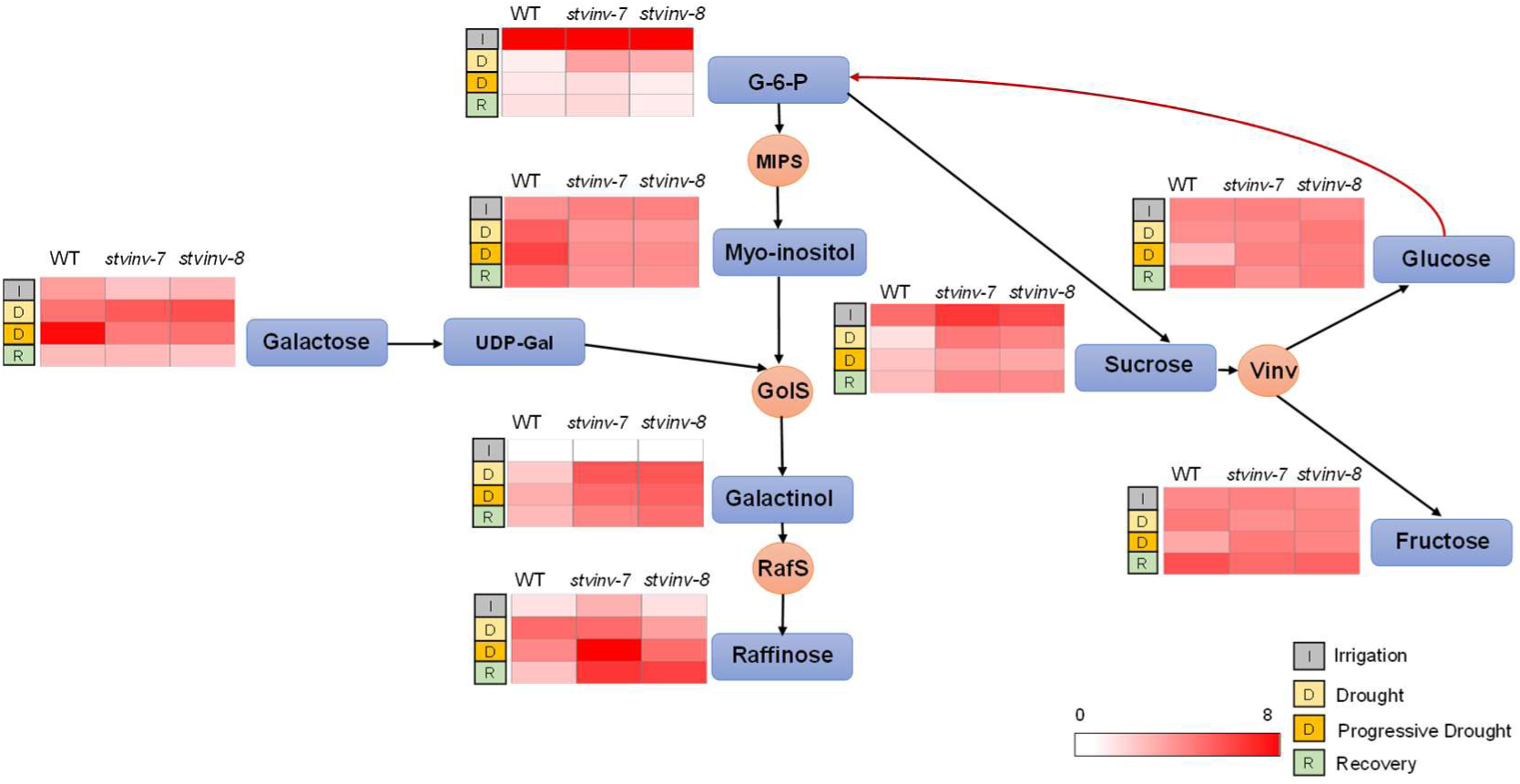
Schematic overview of raffinose family oligosaccharide (RFO) metabolism in WT and *stvinv* plants across irrigation, drought, and recovery phases. The diagram illustrates key metabolites involved in the RFO pathway, including glucose-6-phosphate (G-6-P), glucose, fructose, sucrose, galactose, UDP-galactose, myo-inositol, galactinol, and raffinose. Metabolic conversions are indicated by arrows and associated enzymes: myo-inositol phosphate synthase (MIPS), galactinol synthase (GolS), raffinose synthase (RafS), and vacuolar invertase (VInv). Adjacent heatmaps represent the relative abundance of each metabolite in WT, *stvinv-7*, and *stvinv-8* under irrigation (I), drought (D), and recovery (R) conditions (darker red = higher abundance).

### Drought tolerance of *stvinv* plants is associated with reduced ABA and its catabolites

To assess the hormonal response associated with early drought adaptation, ABA and its catabolic derivatives phaseic acid (PA) and dehydrophaseic acid (DPA) were quantified using LC-MS. Samples were collected from one-month-old plants at baseline (prior to drought; irrigation phase) and after 7 days of drought exposure. As expected, all genotypes exhibited elevated ABA levels in response to drought.

However, *stvinv-7* and *stvinv-8* accumulated significantly lower levels of ABA, PA, and DPA during drought, compared to WT plants (Fig. 7). These findings suggest that drought tolerance in *stvinv* plants may be facilitated by controlled ABA response, complimented by induction of higher RFO levels.

**Fig. 7.**
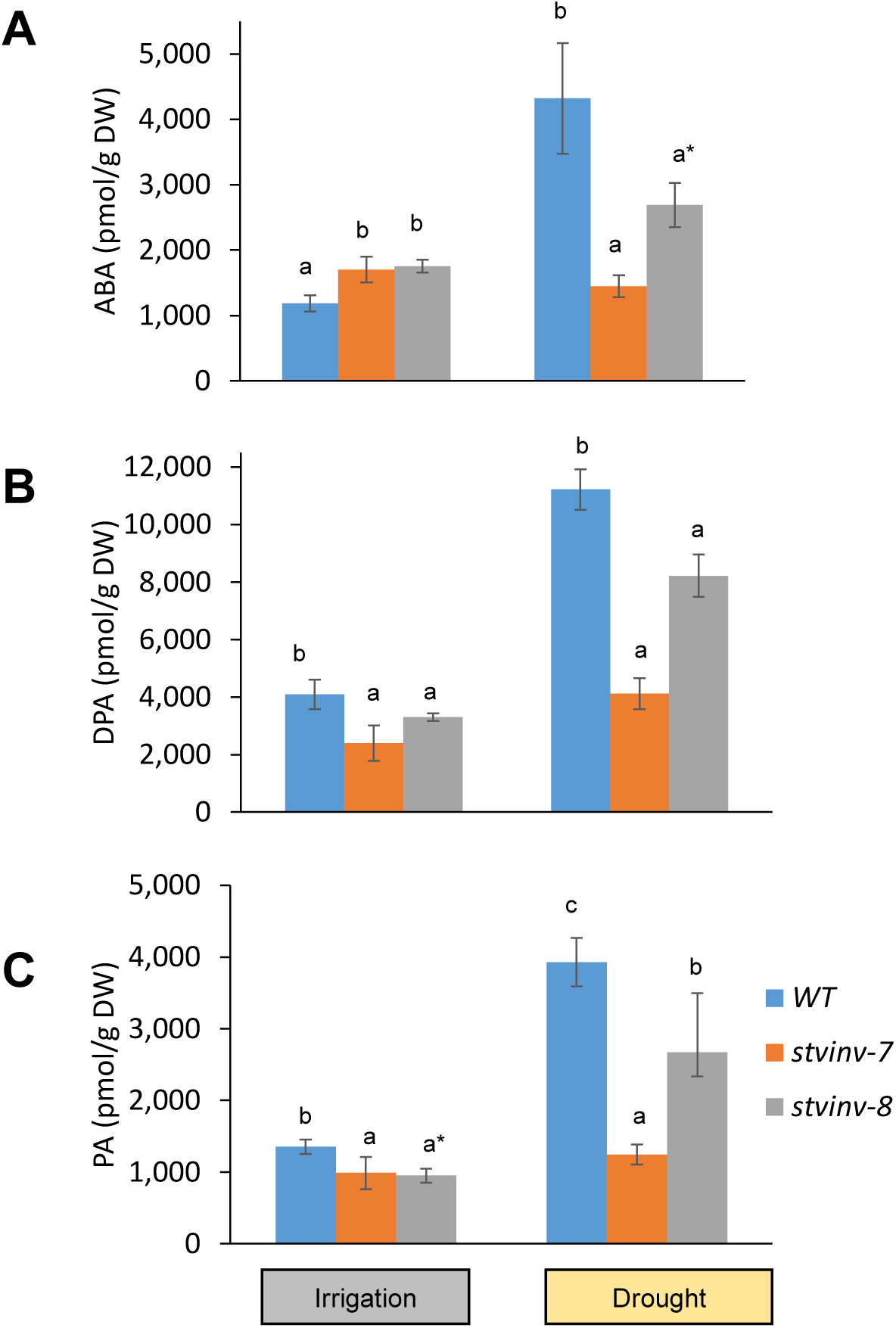
Reduced ABA accumulation and catabolism in *stvinv* plants under drought stress. **A–C,** Concentrations of abscisic acid (ABA), phaseic acid (PA), and dehydrophaseic acid (DPA) were quantified in WT, *stvinv-7*, and *stvinv-8* plants under irrigation and after 7 days of drought using LC-MS. Hormone levels are presented as pmol/g dry weight (DW). Data represent means ± SE (*n* = 5–6 plants per group). Different letters denote statistically significant differences between genotypes within each treatment (Student’s *t*-test, *p* < 0.05); asterisks (*) indicate marginal significance (*p* = 0.05–0.1).

## Discussion

### Enhanced drought resilience and resource efficiency in *stvinv* plants

Drought is a major constraint on potato production, and its impact is expected to worsen with climate change [45]. It has been shown to disrupt carbon partitioning, leading to sugar accumulation in sink leaves and reduced tuber yield and number [45, 46]. The *stvinv* plants exhibited significantly enhanced drought resistance by maintaining consistently higher stomatal conductance, transpiration rates, and photosynthetic efficiency compared to WT plants during both drought and recovery phases (Figs. 1 and 2). Typically, drought stress induces stomatal closure to reduce water loss, but this response comes at the cost of CO₂ assimilation and metabolic activity [59–61]. In contrast, *stvinv* plants maintained open stomata under drought, supporting an unconventional (risk taking) strategy that enables continuous carbon assimilation and sustained energy production despite water limitations. A key advantage of this strategy lies in the knockout plants’ ability to balance elevated transpiration with efficient internal water regulation and minimal tissue damage as revealed by its fast recovery. Despite higher transpiration rate, *stvinv* plants displayed superior AWUE, likely facilitated by their enhanced osmotic adjustment and anticipatory metabolic responses (Fig. 3). Metabolomic profiling of *stvinv* and WT plants revealed significant accumulation of osmoprotective sugars, particularly galactinol and raffinose (Fig. 5), which help maintain cellular water balance and preserve turgor under drought [19, 62]. In addition to supporting photosynthesis, elevated transpiration rate in *stvinv* plants increases evaporative cooling, effectively lowering leaf temperature [63], which in turn, curtail heat-induced ROS production [60, 64]. These protective effects are further reinforced by the ROS-scavenging function of RFOs [19, 65, 66]. We previously showed that, under cold stress, RFO accumulation in *stvinv* plants was linked to decreased ROS levels [67], suggesting the presence of a conserved sugar-mediated mechanism for mitigating oxidative stress that may also operate under drought conditions.

Recent evidence also indicates that osmolyte accumulation, such as elevated RFOs, may positively influence mesophyll conductance (gₘ), the diffusion of CO_2_ from intercellular spaces to chloroplasts. Enhanced gₘ under drought conditions has been associated with improved photosynthetic performance and water-use efficiency due to better membrane stability and reduced oxidative stress [19, 60, 68–70]. Although gₘ was not directly measured in our study, the observed physiological and metabolic traits of *stvinv* plants, including sustained gas exchange, enhanced osmolyte accumulation, and stable photosynthetic capacity (Figs 1-3) are consistent with a scenario in which improved gₘ contributes substantially to their drought resilience.

*stvinv* plants maintain both elevated transpiration and biomass accumulation (Figs 1-4), demonstrating resilience to drought stress. Importantly, this increase in biomass accumulation, particularly in shoot tissue, is not accompanied by a reduction in tuber productivity (Fig. 4). While total tuber weight did not significantly differ between genotypes, *stvinv-*7 produced a slightly higher number of tubers under drought (Fig. 4). This finding suggests that the enhanced shoot vigor in *stvinv* plants, reinforcing their agronomic potential in drought-prone environments allowing the plant to survive longer and potentially produce higher yield.

The rapid recovery of *stvinv* plants following rehydration (Fig. 2) suggests an enhanced capacity to maintain or quickly restore metabolic function. Such resilience could be mediated by sustained sugar signaling and a reduced internal perception of stress, potentially involving transcriptional or epigenetic memory mechanisms [71, 72]. Indeed, the muted accumulation of drought-induced stress metabolites such as proline, homoserine, and shikimic acid in *stvinv* plants (Figs 5 and S2) supports the hypothesis that these plants inherently perceive lower stress levels under drought compared to WT plants.

Collectively, these findings indicate that *stvinv* plants adopt a productive drought strategy, integrating elevated transpiration, osmolyte-based stress buffering, evaporative cooling, and potentially primed recovery mechanisms to sustain growth and yield.

### Resilience supported by metabolic shifts and RFO pathway induction

The drought resilience of *stvinv* plants is further supported by extensive metabolic reprogramming. Elevated levels of citric acid and enhanced activation of glyoxylate and dicarboxylate metabolism pathways indicate improved carbon recycling and energy conservation under stress [73]. These changes likely provide sufficient ATP to sustain basal metabolism and enable rapid recovery after rehydration.

A central aspect of this metabolic shift is the robust activation of the RFO biosynthetic pathway. Metabolomic profiling revealed elevated galactinol and raffinose levels in *stvinv* plants under drought, along with reduced myo-inositol content, indicating a preferential flux toward RFO production. These sugars are known osmoprotectants that stabilize membranes and proteins, helping maintain turgor and mitigate dehydration effects [74–76].

In addition to osmoprotection, galactinol and raffinose act as direct ROS scavengers, contributing to oxidative stress reduction [65, 77–79]. Their accumulation likely reduces reliance on energy-intensive antioxidant systems. This is consistent with the reduced accumulation of stress markers like proline, homoserine, and shikimic acid in *stvinv* plants. Our previous work showed that under cold stress, elevated RFO levels in *stvinv* plants were associated with reduced ROS accumulation [67], suggesting that similar sugar-based mechanisms operate under drought.

The enhanced expression of *GolS3* and *MIP2*, key genes in galactinol biosynthesis and sugar transport, further supports the notion that *stvinv* plants prioritize RFO metabolism under stress. High sucrose levels detected in *stvinv* plants may serve not only as a carbon source for RFO synthesis but also as part of an alternative metabolic route, as proposed in our previous work [67]. Under cold stress, *stvinv* plants shifted toward RFO-producing pathway in the absence of vacuolar invertase, a mechanism likely conserved under drought. Recent findings also suggest that ABA signaling, via ABF-type transcription factors such as CsABF8, can activate RFO biosynthetic genes under drought [80]. The ability of *stvinv* plants to maintain high RFO levels despite low ABA concentrations implies the existence of ABA-independent or sugar-mediated regulatory routes, reinforcing the functional redundancy and plasticity of RFO-inducing mechanisms under stress.

As a central soluble carbohydrate, sucrose may support drought resilience by serving as a rapid energy source, stabilizing osmotic potential, and promoting turgor maintenance [81]. Additionally, sucrose acts as a precursor for RFO biosynthesis and a signaling molecule that can attenuate stress-induced ABA responses or modulate gene expression through ABA-independent pathways [77, 82–86].

Although direct ROS scavenging by sucrose is less established than for RFOs, it may contribute modestly to redox buffering via sugar-regulated ROS genes and interactions with other antioxidants [66, 81, 86–88]. Its primary impact, however, likely reflects energetic efficiency and signaling functions. Together, these roles support the view that elevated sucrose in *stvinv* plants enhances metabolic flexibility and hormonal balance under drought, while promoting the biosynthesis of protective oligosaccharides.

This sugar-centric metabolic strategy may also reflect a broader shift from nitrogen-intensive stress responses (e.g., proline accumulation) to carbon-based protective mechanisms (Figs. 5 and 6). This shift could reduce the metabolic cost of stress adaptation, preserving nitrogen for growth-related functions [89–91]. This is further supported by the consistently lower levels of classical stress markers, indicating that *stvinv* plants likely perceive and experience less physiological stress compared to WT under drought conditions.

Together, the coordinated accumulation of protective sugars, reduced ROS burden, energy-saving metabolic shifts, and lower stress metabolite levels define a sugar-centered, anticipatory strategy that buffers stress perception and sustains physiological function under drought. This profile resembles anisohydric “risk-taking” behavior—maintaining stomatal opening and productivity under moderate drought—and relies on tightly linked hormonal control, in which attenuated ABA signaling and sugar-mediated modulation jointly contribute to drought resilience [92].

### Attenuated ABA signaling and sugar-mediated hormonal modulation contribute to drought resilience

ABA, which is synthesized mainly in the leaves [93], is a central regulator of plant drought responses, typically inducing stomatal closure and restricting growth to conserve water [94–96]. However, *stvinv* plants accumulated significantly lower levels of ABA and its catabolites (PA and DPA) under drought compared to WT plants (Fig. 7). This attenuated ABA response likely reduces the growth penalties commonly associated with prolonged ABA signaling [97, 98], enabling *stvinv* plants to maintain higher stomatal conductance and thus sustain gas exchange, carbon assimilation, and growth even under limited water availability.

A probable mechanism behind this hormonal modulation involves sugar signaling, particularly via hexokinase (HXK), which functions both as a glycolytic enzyme and a glucose sensor modulating ABA responses [82, 99]. Reduced vacuolar invertase activity in *stvinv* plants could limit cytosolic glucose availability, dampening HXK-mediated ABA signaling and facilitating prolonged stomatal opening, thus maintaining physiological activity during drought. This risk-taking strategy could potentially expose plants to desiccation, xylem embolism, or other hydraulic damages. However, the rapid post-drought recovery of *stvinv* plants suggests that they did not incur significant costs for sustaining higher activity under water limitation. It is possible that this resilience is linked to their distinct metabolic adjustments, which mitigate stress damage and support continued function under adverse conditions. The metabolic profile of *stvinv* plants further supports this model. Unlike WT plants, which accumulate stress metabolites such as proline, homoserine, and shikimic acid, *stvinv* plants exhibited reduced levels of these compounds, highlighting their inherently lower stress perception and reduced internal stress signaling [100]. Instead, their metabolic strategy favored accumulating raffinose and galactinol, which provide robust osmoprotection and ROS detoxification, contributing to their drought resilience [36, 74, 79, 101]. Moreover, our previous research under cold stress demonstrated a clear correlation between enhanced RFO accumulation and reduced ROS levels in *stvinv* plants [29]. Given the comparable metabolic response observed under drought, it is plausible that sugar-mediated hormonal crosstalk, potentially involving HXK, ABA, and RFO metabolism, constitutes a conserved regulatory module that buffers stress perception, supports physiological homeostasis, and improves resilience across various abiotic stresses.

Recent studies have further highlighted the role of sugars in modulating ABA sensitivity and response through sugar-responsive transcription factors and signaling pathways, reinforcing the importance of sugar-ABA crosstalk in environmental stress tolerance [102–104]. Such integration likely contributes significantly to the *stvinv* plants’ efficient stress response, energy conservation, and sustained growth.

This shift from reactive to buffered responses may enable *stvinv* plants to avoid costly metabolic penalties associated with conventional stress responses, thus defining a novel and energy-efficient adaptation strategy under drought conditions.

### Non-structural stomatal regulation complements metabolic adaptation

Despite their elevated stomatal conductance, *stvinv* plants did not differ from WT in stomatal density or aperture size. This observation highlights that drought resilience in *stvinv* plants does not result from structural leaf adaptations but rather from functional plasticity in stomatal regulation. Such functional plasticity, modulated by sugar and ABA signaling pathways, allows dynamic optimization of water loss and carbon assimilation under varying environmental conditions (Fig. 8) [59, 105].

**Fig. 8.**
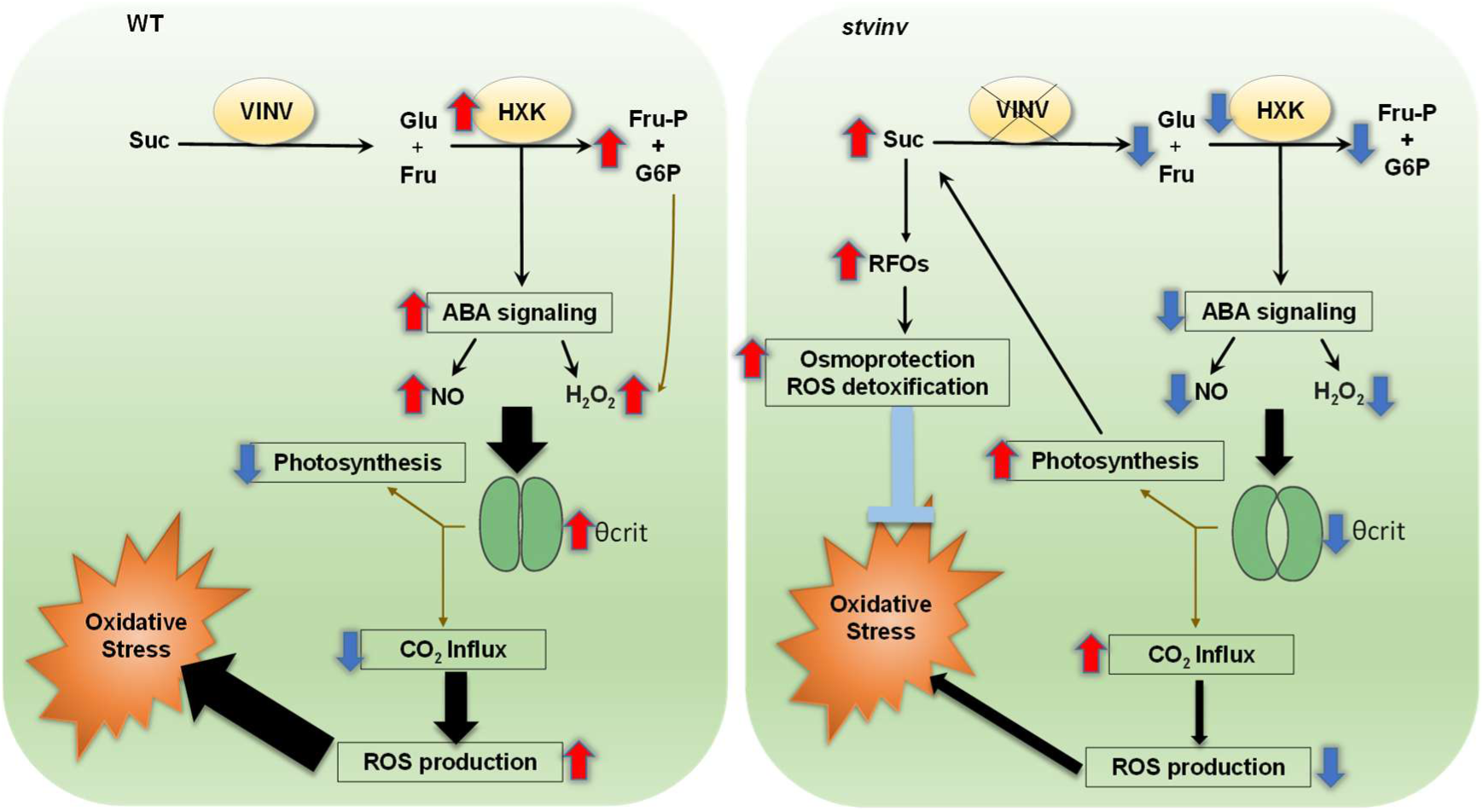
Proposed mechanistic model for enhanced drought tolerance in *stvinv* plants. Schematic model illustrating the differential metabolic and signaling responses to drought stress in WT and *stvinv* plants. In WT plants (left panel), sucrose is cleaved by vacuolar invertase (VINV) into glucose (Glu) and fructose (Fru). These hexoses are phosphorylated by hexokinase (HXK), leading to the accumulation of fructose-6-phosphate (Fru-P) and glucose-6-phosphate (G6P). HXK-mediated sugar signaling enhances abscisic acid (ABA) signaling and increases the production of nitric oxide (NO) and hydrogen peroxide (H₂O₂). This signaling cascade drives stomatal closure at the critical threshold (θcrit), reduces CO₂ influx, and decreases photosynthetic activity. In parallel, enhanced ROS production amplifies oxidative stress and ultimately compromises drought resilience. In *stvinv* plants (right panel), suppression of vacuolar invertase activity results in sucrose accumulation and enhanced synthesis of raffinose family oligosaccharides (RFOs). These sugars provide osmoprotection and facilitate ROS detoxification, thereby alleviating oxidative stress. Attenuated HXK activity reduces ABA-dependent signaling, leading to lower NO and H₂O₂ levels and consequently a weaker induction of stomatal closure. As a result, CO₂ influx is maintained, photosynthesis remains higher, and ROS production is suppressed. Together, these responses underlie the superior drought tolerance of *stvinv* plants. In the diagram, red upward arrows indicate increases, blue downward arrows indicate decreases, blue T-shaped bars denote inhibitory effects, thick black arrows highlight major stress-related fluxes toward ROS generation and oxidative damage, thin arrows represent direct metabolic and regulatory connections, and brown arrows indicate indirect regulatory pathways such as HXK-mediated effects on ABA signaling.

Together, these coordinated physiological and metabolic adjustments allow *stvinv* plants to sustain growth and photosynthesis during water deficit. Their drought response strategy represents a transition from passive resistance to active stress management, highlighting their potential as valuable models for engineering stress resilience in crop species.

## Methods

### Plant growth conditions and drought stress treatment

*Solanum tuberosum* cv. Désirée plants were propagated in tissue culture for 3 weeks and transferred to small pots (4.5 × 8.5 cm) containing standard potting mix (Pelemix Green) for an additional 4 weeks. Seedlings were maintained in a controlled growth chamber under a 16 h light/8 h dark photoperiod at 23 °C, with ∼200 µmol m⁻² s⁻¹ photosynthetically active radiation provided by LED illumination.

Plants were irrigated twice daily until stress induction. Drought stress was imposed by withholding irrigation for 15-18 consecutive days, followed by rewatering to monitor recovery responses over a 12-day period. Drought response was visually evaluated using a phenotypic scale ranging from 1 to 10, where a score of 10 represented a high-vigor plant and a score of 1 indicated severe wilting. Physiological and biochemical measurements were conducted before, during, and after the drought cycle to assess plant performance and stress resilience. Each genotype was represented by a minimum of 8 biological replicates (8–12 plants per treatment), ensuring robust statistical evaluation of drought responses.

### Physiological measurements under drought stress

Leaf stomatal conductance and chlorophyll fluorescence were measured using the LI-600 and LI-600N Porometer/Fluorometer (LI-COR Biosciences), which simultaneously records gas exchange and photosynthetic efficiency from the same leaf area. Measurements were performed daily on fully expanded mature leaves from the second node below the apical bud, at a fixed time point, one hour after lights-on. This time was chosen to allow for stabilization of photosynthetic activity following the dark-to-light transition, while minimizing diurnal variability.

Chlorophyll content was assessed non-destructively using a SPAD-502 Plus Chlorophyll Meter (Konica Minolta, Japan), which estimates relative chlorophyll concentration based on leaf light transmittance. Measurements were performed on the same leaves used for gas exchange analyses, with three readings per leaf averaged to yield a single value per plant. At least 10 biological replicates per genotype were analyzed across all time points.

### Whole plant water relations phenotyping using a lysimetric array (Plantarray)

Physiological phenotyping, including whole-plant measurement of transpiration dynamics, canopy stomatal conductance, and water-use efficiency (WUE), was performed using the Plantarray 3.0 high-throughput functional phenotyping platform (Plant-DiTech, Israel), following established protocols [106]. Potato tuber sprouts were excised at the nodal region and planted in a soil mixture consisting of polystyrene: vermiculite: peat moss in a 3:2:1 ratio (v/v). Seedlings were grown in a greenhouse under natural day length and ambient Mediterranean winter conditions; temperatures ranging between 15°C and 32°C, relative humidity between 25% and 40%, and a photoperiod of ∼11 h light/13 h dark, with all environmental parameters (temperature, humidity, PAR, VPD) continuously monitored, see supplementary Fig. S4 [106, 107]. Average PAR during daytime was ∼600–1200 µmol m⁻² s⁻¹. After one month, plants were transferred to the ICORE Center for Functional Phenotyping greenhouse for an additional month of acclimation under daily irrigation. Subsequently, two-month-old plants were transferred to the lysimetric array and grown for 34 days (3 March–4 April 2022) in 4-L pots filled with sand 20/30 (Negev Minerals) under natural sunlight and moderately controlled temperature conditions, simulating near-field environments [108, 109]. To prevent soil evaporation, pots were covered with a plastic film.

The experiment was conducted in a randomized block design. The experiment followed a randomized design, with 8–14 biological replicates per genotype in the drought treatment and 3–4 well-watered controls per genotype maintained under full irrigation. Soil volumetric water content (VWC) was continuously monitored via 5TE (Meter, USA) sensors placed in each pot, while environmental variables were recorded to compute vapor pressure deficit (VPD: 0.7–4.0 kPa) (Supplementary Fig. S4). Whole-plant transpiration-rate was measured gravimetrically at 3-minute intervals by analyzing changes in pot weight using high-resolution load cells, enabling continuous monitoring of water fluxes in the soil-plant-atmosphere continuum [108]. A three-phase irrigation regime was applied: (i) well-watered pretreatment (days 1–12); (ii) drought induction (days 13–27), during which irrigation was reduced daily to 80% of each plant’s previous-day transpiration, mimicking progressive soil water depletion as occurs in field conditions [110]; and (iii) full re-irrigation recovery (days 28–34). The Plantarray platform includes a feedback-controlled irrigation system that ensures uniform drought exposure across genotypes by tailoring water delivery to each plant’s individual transpiration rate [106]. Processed data were analyzed via the SPAC Analytics web tool, which supports both real-time visualization and statistical analysis of multiple parameters, including transpiration rate, stomatal conductance, growth rate, and WUE. WUE was calculated daily as the ratio of modeled daily biomass gain to daily transpiration, as implemented in SPAC Analytics. Agronomic water-use efficiency (AWUE) was computed per plant as final total dry biomass divided by cumulative transpiration over the full experimental period: AWUE = (DW_shoot + DW_tuber + DW_root) / ∫Transpiration dt, where dry biomass was determined at harvest after oven-drying, and cumulative transpiration (L plant⁻¹) was obtained by time-integrating the load-cell mass balance with irrigation inputs and drainage subtracted; units g L⁻¹.

### Tissue sampling

For metabolomic analysis, fully expanded young leaves were sampled from the third node below the apical bud at six distinct time points: T0 (baseline, following 10 days of full irrigation), T1 and T2 (mild drought, after 7 and 9 days without irrigation, respectively), T3 and T4 (progressive drought, after 12 and 14 days without irrigation), and T5 (recovery, one day after rewatering). For ABA quantification, leaf samples were collected at two time points: baseline (during the irrigation phase) and at the middle of the drought phase (day 7). All samples were collected at midday from well-expanded leaves to minimize diurnal variation, immediately frozen in liquid nitrogen, and stored at –80 °C until further analysis.

### Quantitative analysis of ABA metabolites

Endogenous levels of ABA and its metabolites were assessed as previously described: sample purification and chromatographic separation was performed according to Danieli et al. [111] and settings for the MS detection were according to Vrobel et al. [112]. Briefly, 3 mg of homogenized lyophilized leaves were extracted with 1 ml of 50% aqueous acetonitrile containing a mixture of internal standards (4 pmol of D6-ABA, D3-PA and D3-DPA per sample) on ice-cold ultrasonic bath for 30 min. After centrifugation (20 000 g, 10 min, 4°C) the supernatant was loaded onto an Oasis HLB column (1 ml cartridge, 30 mg sorbent, Waters, USA) equilibrated with 1 ml methanol, 1 ml water and lastly by 1 ml 50 % acetonitrile. Flow-through fraction was collected and pooled with elution solvent, 1 ml of 30% acetonitrile. Pooled fractions were evaporated *in vacuo*. Prior to analysis, samples were resuspended in 40 µL of mobile phase and analyzed. Analyses were performed using a Nexera X2 modular liquid chromatograph coupled to an MS 8050 triple quadrupole mass spectrometer (Shimadzu) via an electrospray interface. Chromatographic separation was performed using Waters analytical column CSH™ C18, 2.1 mm × 150 mm, 1.7 µm. Aqueous solvent A consisted of 15 mM formic acid adjusted to pH 3.0 with ammonium hydroxide. Solvent B was acetonitrile. Separation was achieved by gradient elution at a flow rate of 0.4 mL/min at 40°C: 0–1 min 20% B; 1–11 min 80% B in a linear gradient, followed by washing and equilibration to initial conditions for a further 7 min. Three MRM transitions were monitored for each analyte (ABA, PA, DPA, ABA-GE and internal standards) to ensure correct identification. Raw data were processed using Shimadzu software LabSolutions ver. 5.97 SP1.

### Extraction of primary metabolites

Metabolite analysis by GC-MS was carried out by a method modified from that described previously (Roessner et al., 2001) extracted in 1 ml prechilled methanol:chloroform:water extraction solution (2.5:1:1 v/v) with 380 µl of Standards (1 mg/ml ribitol in water) subsequently added as a internal standard. The mixture was sonication for 10 min, shaking for 10 min at 25°C. After centrifugation at 14000 RPM, 300 µL of water and chloroform was added to the supernatant. Following vortexing and centrifugation the methanol-water phase was taken and kept at −80°C until use.

### Derivatization and analysis of primary metabolites in GC-MS

200 µL of methanol-water phase reduced to dryness in vacuum. Residues were redissolved and derivatized for 120 min at 37°C in 40 µL of 20 mg/mL methoxyamine hydrochloride in pyridine) followed by a 30-min treatment with 70 µL N-methyl-N-(trimethylsilyl)trifluoroacetamide at 37°C. 7 microliters of a retention time standard mixture (0.029% v/v n-dodecane, n-pentadecane, n-nonadecane, n-docosane, n-octacosane, n-dotracontane, and n-hexatriacontane dissolved in pyridine) was added prior to trimethylsilylation. The first run was done by injecting 1 µL to analyse melibiose and second run was performed with 0.2 µL to analyse galactose. Both runs were done in splitless mode. The GC-MS system consisted of a 7693 autosampler, a 7890B GC, and a 5977B single quadrupole mass spectrometer (Agilent ltd). The mass spectrometer was tuned according to the manufacturer’s recommendations using tris-(perfluorobutyl)-amine (CF43). GC was performed on a 30 m VF-5ms column with 0.25 mm i.d. and 0.25 m film thickness +10 m EZ-Guard (Agilent). (Split/splitless liner with Wool, Restek, USA). Gradient of Injection temperature (PTV) was from 70°C to 300°C in 14.5°C/sec, the Transfer line was 350°C, and the ion source adjusted to 250°C. Gain factor 15. The carrier gas used was helium set at a constant flow rate of 1 ml/ min. The temperature program was 1 min isothermal heating at 70°C, followed by a 1°C/min oven temperature ramp to 76°C, followed by a 6°C/min oven temperature ramp to 340°C, and a final 5 min heating at 340°C. Mass spectra were recorded at 1.6 scans per second with a mass-to-charge ratio 70 to 550 scanning range. Post-column Back-flushing was used during the post-run time in every injection to keep detector clean, column flow was reversed for few minutes to remove high-boiling components to inlet split vent. JetClean procedure was used after 1 µL set (15 injections) to keep the ion source clean, based on reductive hydrogen cleaning principle. Spectral searching in the Masshunter software (Qualitative and Unknown analyses) against RI libraries downloadable from the Max-Planck Institute for Plant Physiology in Golm, (http://gmd.mpimp-golm.mpg.de/) the result normalized by the internal standard ribitol and the average of the pools, and finally by log transformation.

### Metabolomic data analysis

Raw metabolite abundance data were preprocessed, normalized, and statistically analyzed using the web-based platform MetaboAnalyst 5.0 (https://www.metaboanalyst.ca)[113]. Data normalization included log transformation and auto-scaling to facilitate downstream multivariate analyses. Principal component analysis (PCA) and partial least squares-discriminant analysis (PLS-DA) were performed to assess clustering and group separation. Differential metabolites were identified using univariate statistics (Student’s *t*-test, FDR-adjusted *P* < 0.05) and visualized using volcano plots and heatmaps. Enrichment and pathway analyses were conducted using the *Arabidopsis thaliana* reference pathway library, based on KEGG and SMPDB

### Stomatal density and aperture

Stomatal traits were quantified using a rapid dental-resin imprinting method following Geisler [114, 115]. Fully expanded leaves that had reached final size were sampled from the third node below the apical bud, from well-watered plants grown under controlled long-day conditions (16 h light: 8 h dark). Sampling was conducted at mid-morning, approximately 1 h after lights-on (09:30–10:30 local time), to minimize diurnal variation in stomatal aperture. Stomata and impressions of the abaxial (lower) and adaxial (upper) epidermis were taken by gently applying a thin layer of clear dental resin (Zhermack Elite HD). After drying (∼10–15 min), the resin was covered with nail polish and covered with film. The film was peeled off and mounted onto microscope slides. This method preserves stomatal morphology and distribution with high fidelity, allowing for accurate microscopic analysis.

Imprints were visualized using bright-field light microscopy at ×200–×400 magnification. Stomatal density (number of stomata per mm²) was quantified by counting stomata within defined calibrated areas. Stomatal aperture width, defined as the distance between the inner edges of the guard cells, was measured using an eyepiece micrometer and validated by ImageJ, a digital image analysis software [116].

Multiple fields per sample (typically 3–5) were analyzed to account for spatial variability across the leaf surface, and at least three biological replicates per genotype were used. This approach enabled us to detect subtle differences in stomatal morphology and behavior under drought and control conditions.

### Harvest and biomass measurements

At the conclusion of the experiment, whole plants were harvested and separated into three major components: aboveground green tissues (including fully expanded leaves and stems), roots, and tubers. Each plant part was immediately weighed to determine fresh biomass. Leaf and stem tissues were then air-dried at room temperature for 7 days, followed by oven-drying at 60 °C until constant weight. The resulting dry weight of aboveground green tissues was recorded for subsequent analysis.

## Data availability

All data supporting the findings of this study are available within the article and its supplementary information files.

## Supplementary figures

**Fig. S1.**
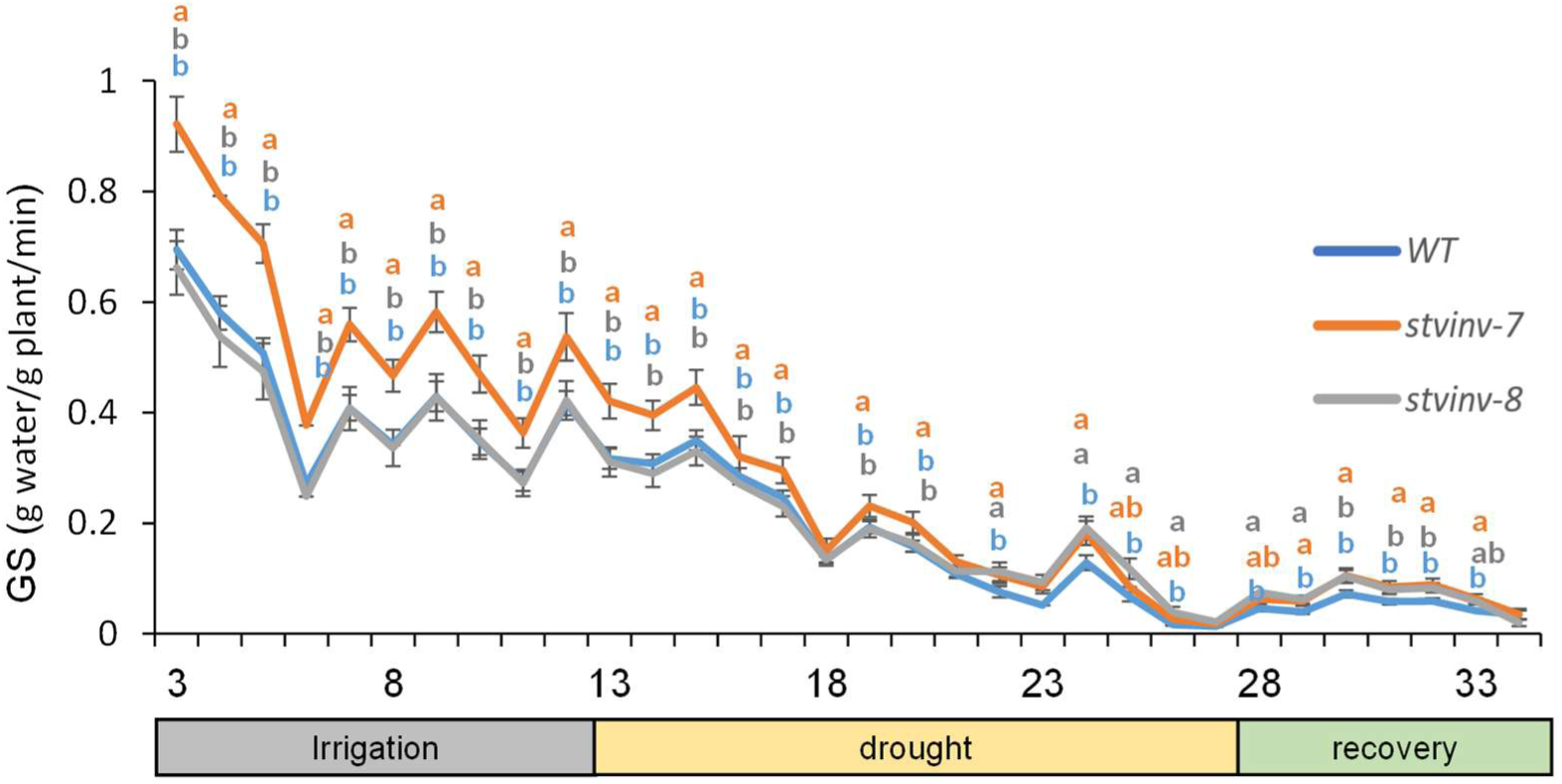
Whole-canopy midday conductance (10:00–15:00). Different letters above data points indicate statistically significant differences between genotypes (p < 0.05).

**Fig. S2.**
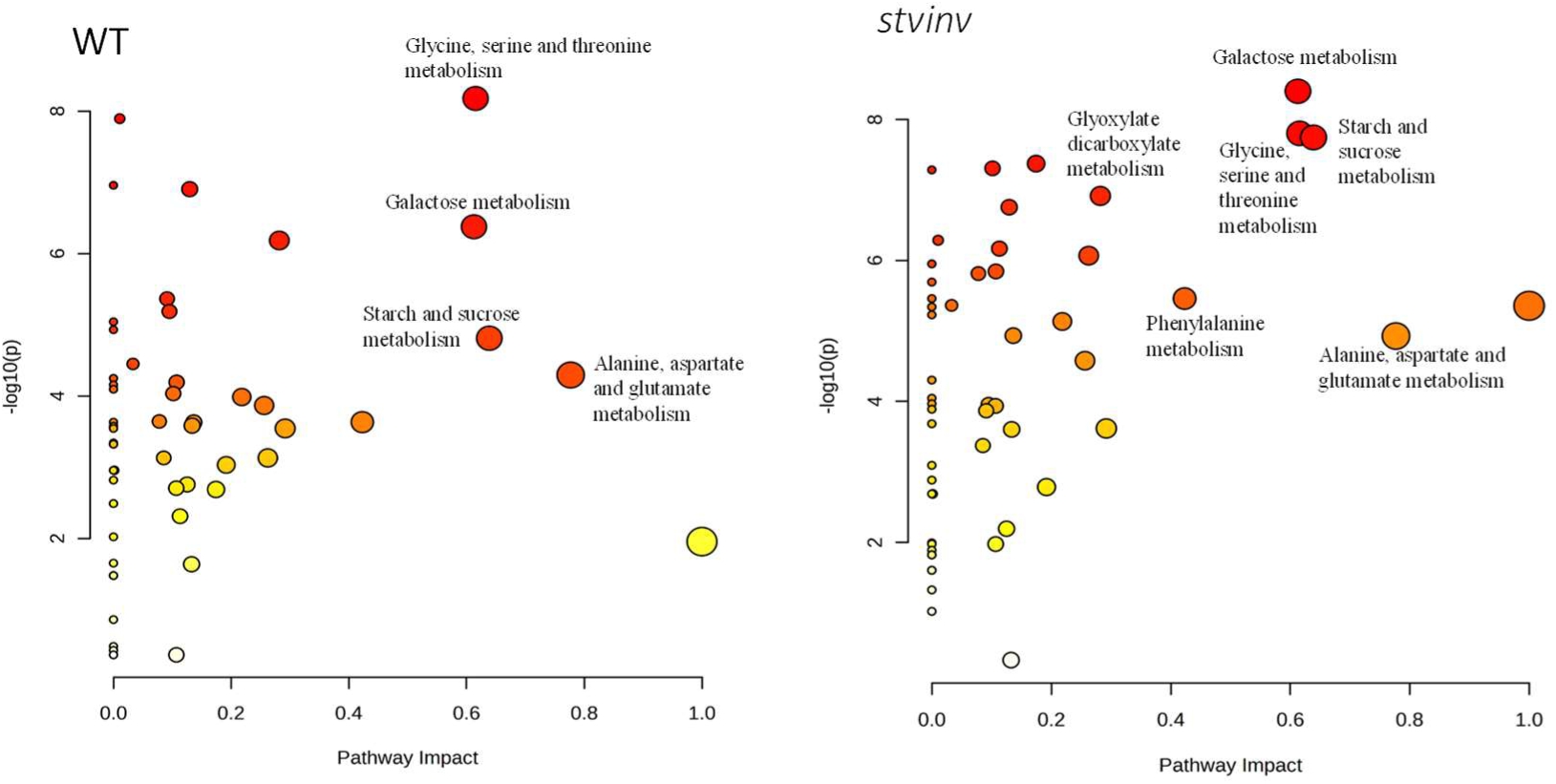
Pathway impact analysis. For WT and *stvinv* plants, with x-axis showing pathway impact (betweenness centrality) and y-axis showing significance (–log₁₀(*p*)).

**Fig. S3.**
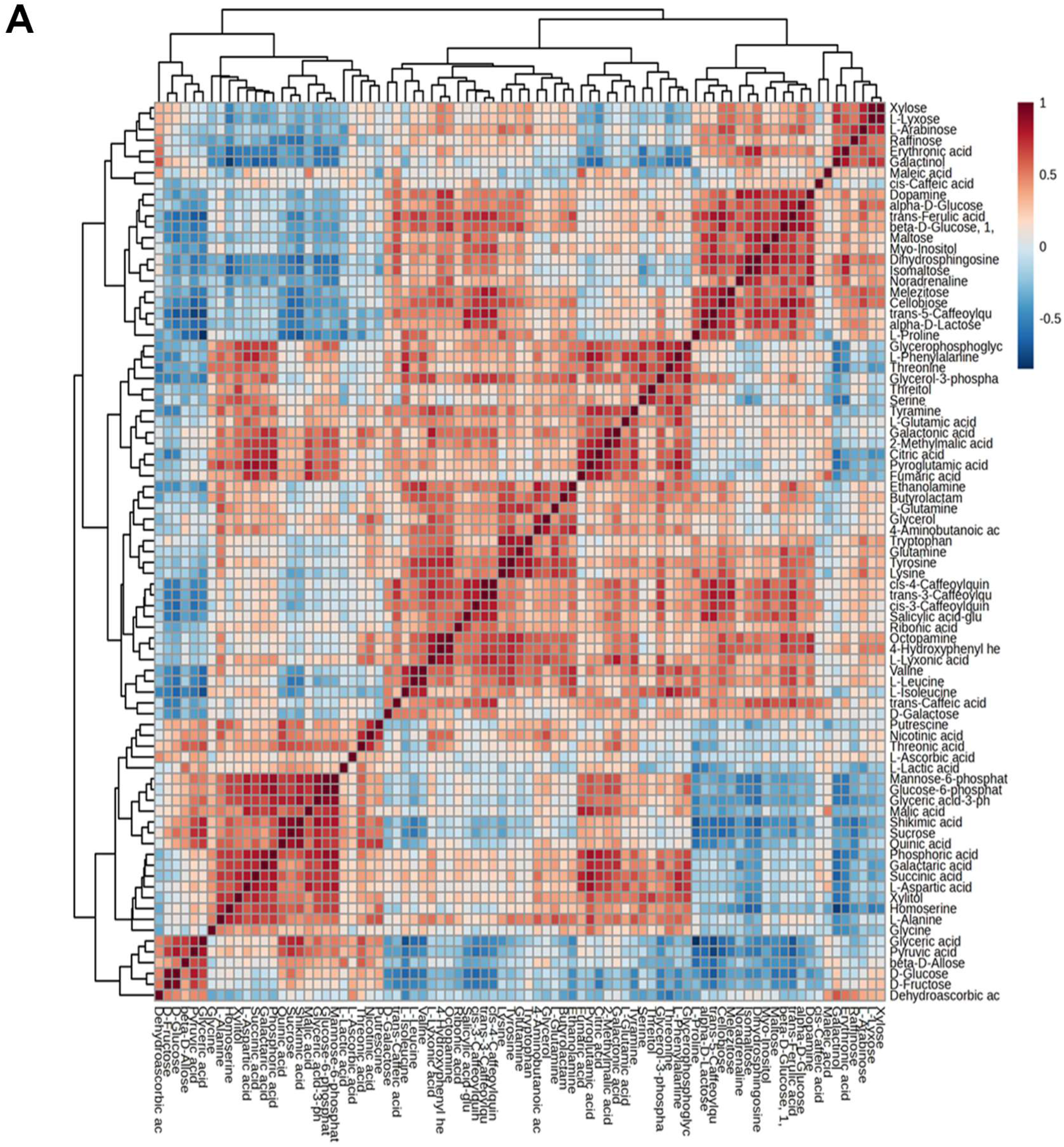
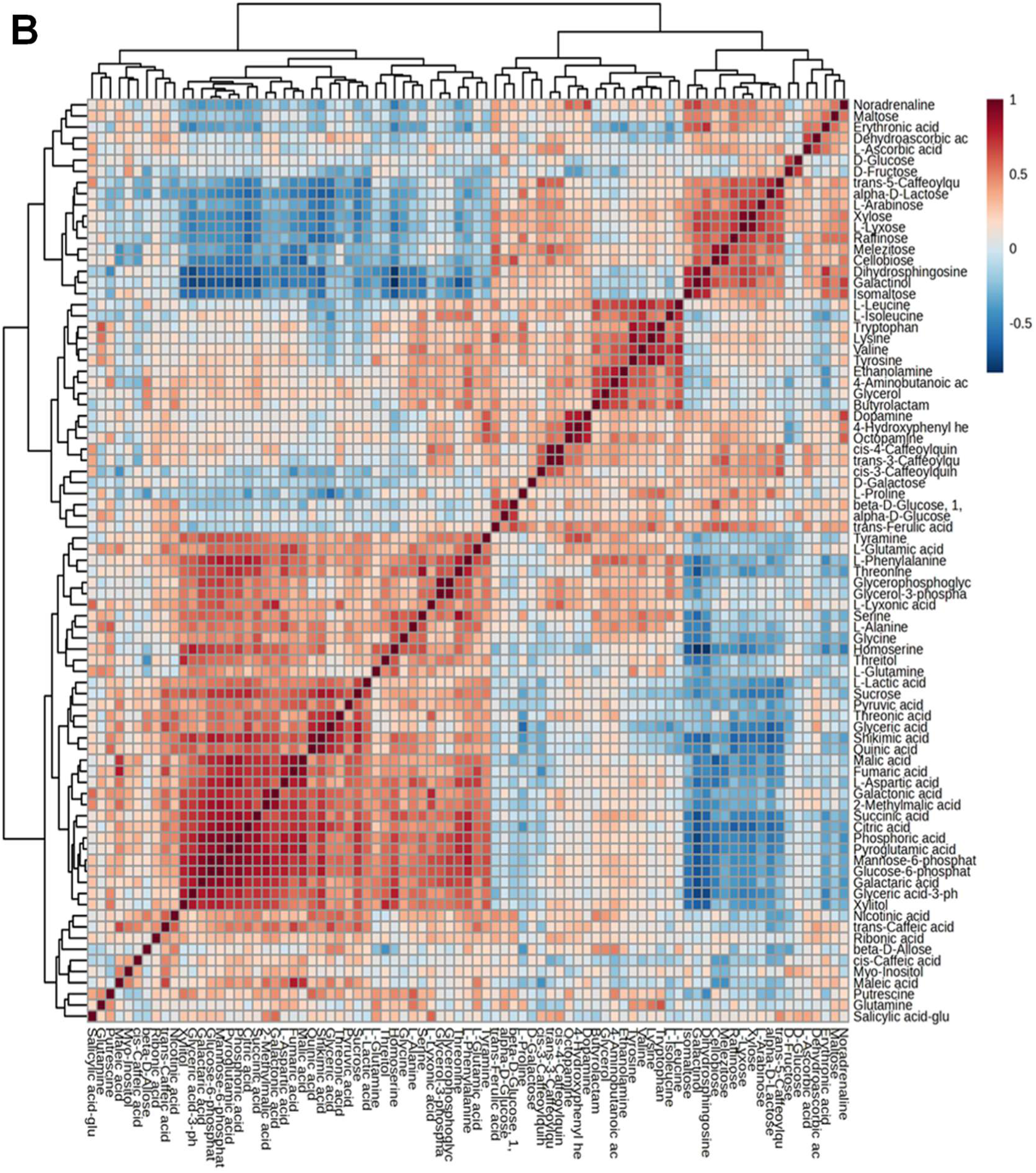
A-B Correlation heatmaps. Reveal metabolic network organization in WT (E) and *stvinv* plants (F). WT plants exhibit broad clustering among carbohydrate, amino acid, and secondary metabolites, reflecting a generalized stress response. In contrast, *stvinv* plants show tight clustering of osmoprotective metabolites, such as sucrose, raffinose, and galactinol, indicating a focused metabolic adaptation to drought.

**Fig. S4.**
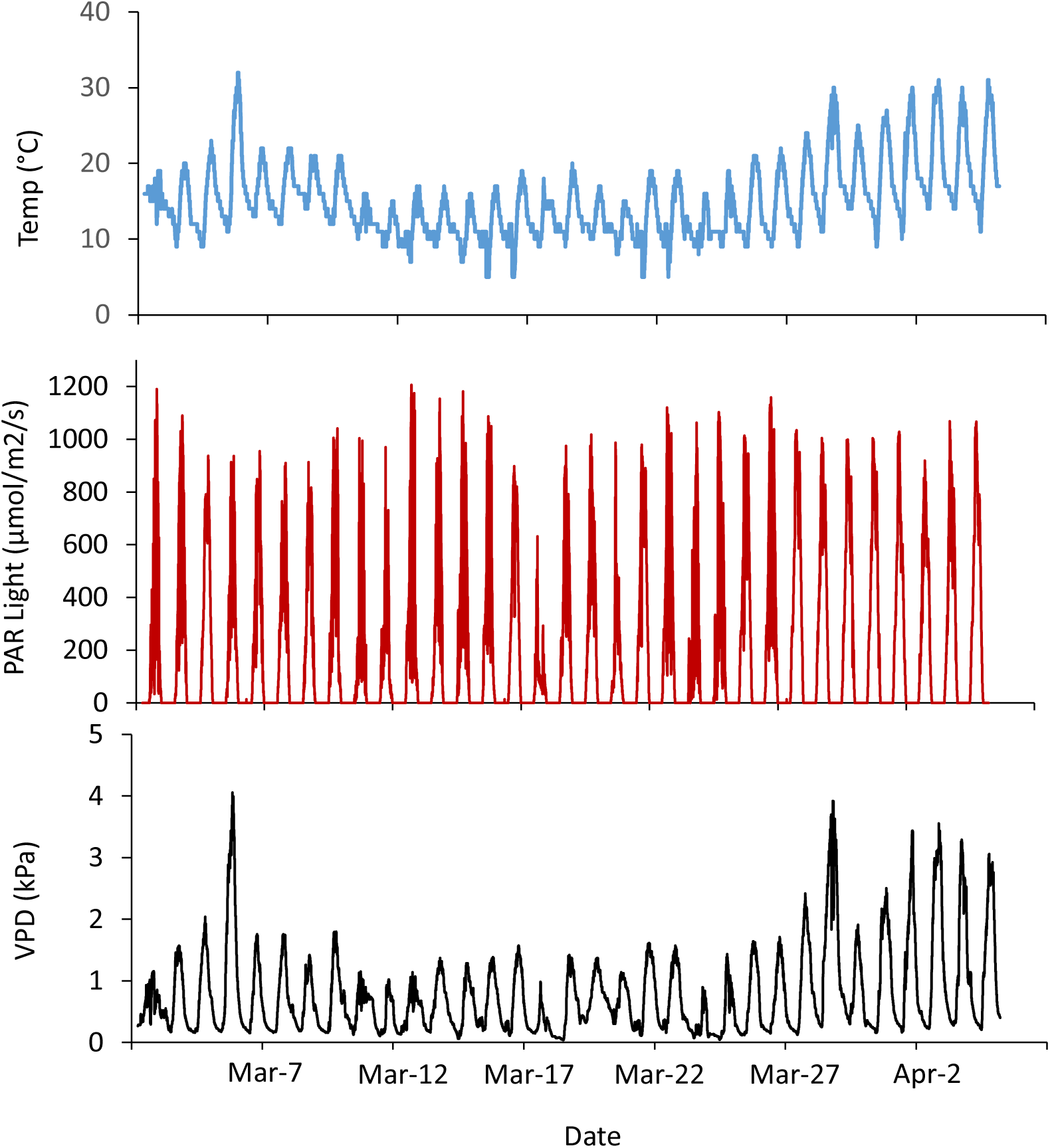
Parameters of the soil–plant–atmosphere continuum monitored by “Plantarray”. (A), air temperature (T, °C). (B), Photosynthetic active radiation (PAR, mmol m-2 s −1). (B), vapor pressure deficit (VPD, kPa). The data shown in the graph were obtained from the spring-season experiment in March - Apil 2022.

## Notes

### Competing Interest Statement

The authors have declared no competing interest.

